# Responses of the *in vitro* turtle brain to visual and auditory stimuli during severe hypoxia

**DOI:** 10.1101/2022.07.01.498485

**Authors:** Michael Ariel, Shivika Ahuja, Daniel E. Warren

**Author notes:** Corresponding Author. Saint Louis University, St. Louis, Missouri 63103 U.S.A.

## Abstract

North American pond turtles (Emydidae) are renowned for their ability to survive extreme hypoxia and anoxia, which enables several species to overwinter in ice-locked, anoxic freshwater ponds and bogs for months. Centrally important for surviving these conditions is a profound metabolic suppression, which enables ATP demands to be met entirely with glycolysis. Despite this, turtles have occasionally been observed exploring their natural and laboratory environments while anoxic and can still respond to sensory stimuli. To better understand whether anoxia limits a special sensory function, we recorded evoked potentials in a reduced brain preparation, *in vitro,* that was perfused with severely hypoxic artificial cerebral spinal fluid (aCSF). For recordings of visual responses, an LED light was flashed onto retinal eyecups while evoked potentials were recorded from the retina or the optic tectum. For recordings of auditory responses, a piezo motor-controlled glass actuator vibrated the tympanic membrane while evoked potentials were recorded from the cochlear nuclei. We found that visual responses decreased even with moderately hypoxic perfusate (aCSF P_O2_ = 30-60 torr) and completely abolished under severe hypoxia. In contrast, the evoked response within the cochlear nuclei was unattenuated with severe hypoxia (aCSF P_O2_ < 20 torr). These data provide further support that pond turtles have a limited ability to sense visual information in their environment even while moderately hypoxic, but that auditory input may become a principal avenue of sensory perception during extreme diving in this species.

**Summary Statement:** Severe hypoxia attenuates the visual, but not auditory responses in a reduced brain preparation from a pond turtle.

## Introduction

Despite the abundance of oxygen in the atmosphere, its availability for aerobic respiratory function can vary considerably for animals, particularly for aquatic species where its solubility in water is low, competition for it can be high, and its exchange with the atmosphere limited. When oxygen levels fall (hypoxia) to undetectable levels (anoxia), survival relies entirely on the ability of that animal to utilize anaerobic pathways for energy production, which, in vertebrates, yields just a fraction of the ATP per mole of glucose compared to aerobic respiration. Consequently, surviving anoxia is highly dependent on an animal’s capacity to decrease its metabolic rate.

Among the champions of vertebrate anoxia-tolerance are several species of North American pond turtle (Emydidae), which can survive anoxia for as many as 5 months at 3°C (Odegard et al. 2018) and for 36 hours at room temperature (Johlin and Moreland 1933). This performance would be impossible without a greater than 70% decrease in metabolic rate, which is achieved through the arresting of protein synthesis, ion channel function, and synaptic transmission (Jackson 2000;(Buck and Pamenter, 2018). In turn, a decreased circulatory requirement further enables suppression of cardiovascular function (Hicks and Farrell, 2000a; Hicks and Farrell, 2000b), producing additional energy savings. Extreme anoxia-tolerance has been exploited by some overwintering pond turtle species (i.e., painted turtles, *Chrysemys picta*) which spend their winters in ice-locked freshwater ponds and bogs that become severely hypoxic and anoxic for many months across their native geographic range.

Although the importance of hypoxia- or anoxia-induced metabolic suppression is without question, the response, itself, produces obvious functional trade-offs for a tissue or organ system. Presumably, those tissues and organs that are either non-essential or energetically expensive during the anoxic episode should be the first to exhibit functional and metabolic suppression, while essential and energetically inexpensive systems will be the last affected. Nervous system function is considered energetically expensive, and predictably exhibits functional suppression, particularly in the cerebrocortex (Buck and Pamenter 2018; Buck et al 2012). Less well-explored are the responses in other regions of the brain, such as the brainstem, where homeostatic pathways might continue to respond to internal physiological signals, as well as external environmental cues during the winter. Indeed, anoxic turtles can be aroused and have been observed exploring their anoxic habitat, which naturally leads to basic questions about the biology of hypoxia-tolerant animals, like turtles: how is the sensory world altered by oxygen deprivation and do those changes affect the way the animal interacts with its environment?

The experiments described herein focus on the well-studied special senses processed in the brainstem that mediate responses to light and sound. External cues, such as photon absorption by retinal photoreceptors and mechanical vibrations of the tympanic membrane, are necessary sensory functions under normal conditions, but their requirements and maintenance during anoxia are likely to differ. Using reduced, *in vitro*, preparations of red-eared slider turtles (*Trachemys scripta elegans*), a common, relatively anoxia-tolerant North American pond turtle, visual and auditory stimuli were presented to neural tissues in either oxygenated or severely hypoxic artificial cerebrospinal fluid. We report that neural responses do, in fact, differ for these two sensory systems during severe hypoxia.

## Materials and Methods

### Animals

Red-ear pond sliders, *Trachemys scripta elegans* (both sexes, 175–250 g, were purchased from Niles Biological, Sacramento, CA), housed in aquaria filled with municipal water (18–22°C) with access to basking platforms illuminated by incandescent bulbs and were fed Aquatic Turtle MonsterDiet (Zeigler Bros., Gardners, PA) thrice weekly, *ad libitum*. All turtle procedures were in strict adherence to the Guide for the Care and Use of Laboratory animals of the National Institutes of Health and approved by the Institutional Animal Care and Use Committee of Saint Louis University (Protocol #2979).

### Preparation of the in vitro brainstem

Following induction of surgical anesthesia (ice for approximately 30 minutes and then an injection of propofol (10-20 mg/kg) into the neck’s sagittal subcarapacial sinus), the head was decapitated and then bathed in ice-cold, oxygenatedaCSF (also referred to as perfusate or media) for subsequent dissection. The composition of the aCSF used throughout the study was always the following (in mM: 96.5 NaCl, 2.6 KCl, 2.0 MgCl_2_, 4.0 CaCl_2_, 31.5 NaHCO_3_; 274 mOsmol). When bubbled with 5% CO, this produced a pH of 7.6 at room temperature. Except when severe hypoxia was imposed, the aCSF was always bubbled with 95% O_2_/5% CO_2_). During hypoxia, the perfused media was simply the control media bubbled with CO_2_ (usually 5%, but in some instances, 3%) with the gaseous balance always being N_2_. There were no differences in responses between the two types of media, so the results from these experiments were combined for analysis and presentation.

The neural tissues were placed in a recording chamber perfused with oxygenated aCSF and fixed to a vibration-isolation table (Vibraplane, Kinetic Systems, Boston, MA). Optodes were placed in the perfusing solution in the chamber (Witrox from Loligo® Systems, Viborg, Denmark) to measure the PO2 continuously, while a recording micropipette was visually guided to its neural target. Three preparations were employed to investigate the two forms of sensory stimulation, as described below.

### Preparation 1: Auditory Recordings from the Cochlear Nuclei

The preparation used to study auditory function involved recording evoked potentials from the cochlear nuclei in an *ex vivo* preparation. Those nuclei are visible in the medial brainstem, after the overlying cerebellum was deflected laterally, revealing them in the cerebellar peduncle adjacent to the vestibular nuclei (Fan et al., 1997). Field potentials were recorded during tympanic membrane (TM) vibration that was then relayed into the temporal bone, transduced by hair cells and transmitted as spike activity to the brainstem via the eighth cranial (vestibulocochlear) nerve.

Drawings and photographs of the TM and the reduced preparation are shown in Figure 1, along with examples of field potential responses recorded in the brainstem. A TM was identified by two concentric wrinkles in the skin on this lateral photograph of a living turtle on the lateral wall of its head, caudal to the eye and mouth, just ventral to its signature patch of red skin (Figure 1C). Figure 1A shows the head structure drawn from above. A probe mounted on a piezo actuator presses on the turtle’s left TM, which is connected via an ossicle (thin rectangle) that transverses in the air-filled middle ear (gray oval) to the transducing organ (otic capsule, a small white oval) in the temporal bone.

**Figure 1.**
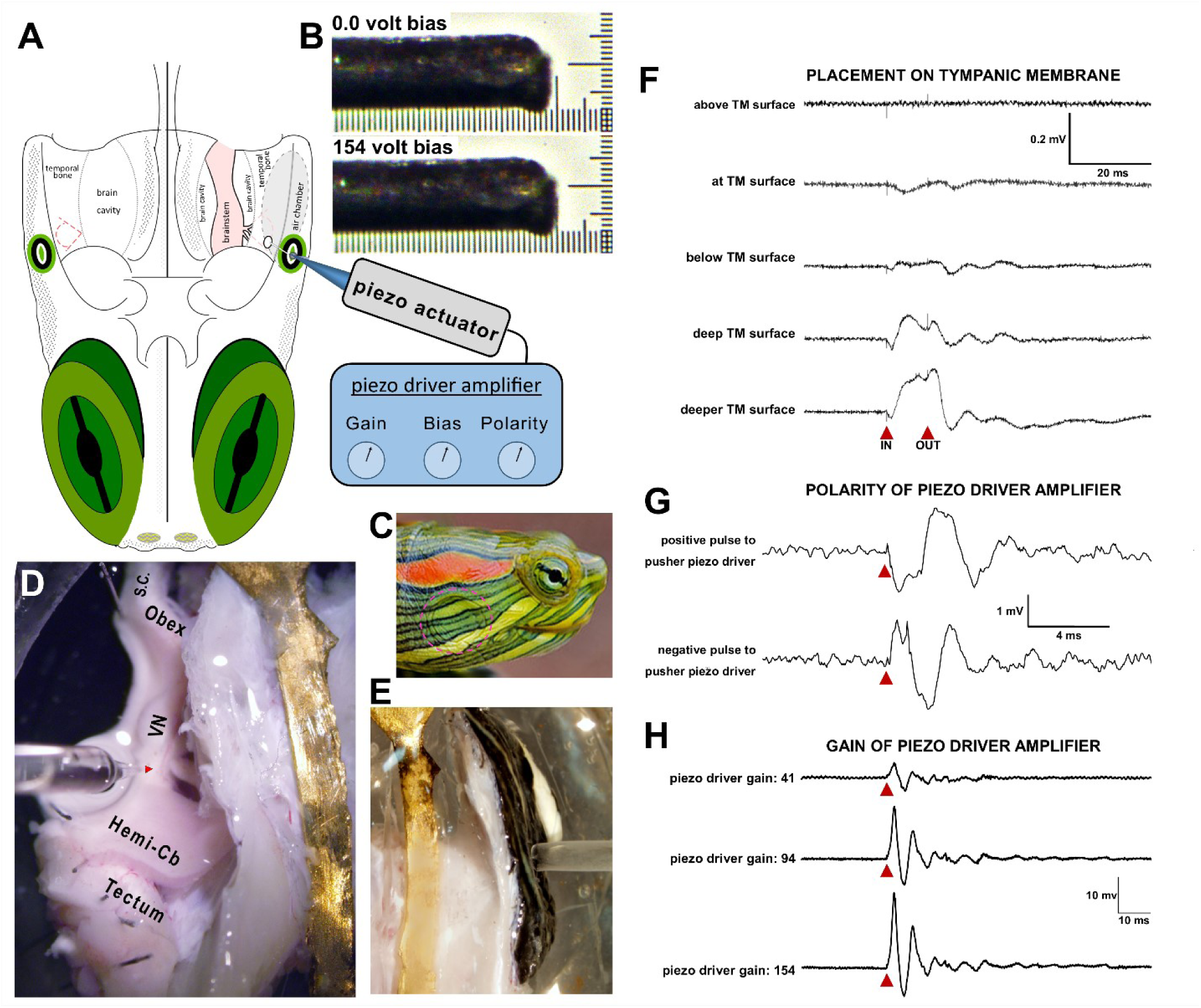
Methods to Move the Tympanic Membrane *in vitro*. **A**) Drawing of dorsal view of pond turtle, nose down, showing the placement of the piezo linear actuator pushing on the left tympanic membrane. Behind the TM, drawn as concentric green/white circles, is a white, thin ossicle that transverses the air chamber to relay TM vibrations to the Otic Capsule within the temporal bone. The two branches of the eighth cranial nerve fuse within the brain cavity and bring sensory signals to the brainstem. **B**) The movement of the piezo pusher was visualized using the dissecting microscope directly above the recording chamber. A digital camera (Olympus DP71) was fitted onto one optical path which also contained a calibration reticle. At the highest magnifications, 10-µm gradations were observed and then photographed at maximal resolution. The position of the piezo pusher was photographed after a thin opaque tungsten rod replaced the glass probe, because the camera did not adequately capture the transparent glass tip. The top image shows the rod’s tip when the piezo stepper was set to the zero-bias volts of the stepper driver (Burleigh PZ150M) which is rated to move from 0-7 µm). The bottom image shows the rod’s tip when the piezo stepper driver was set to its maximum bias level of 154 volts. The difference in position was roughly 7 tenths of the 10-µm gradation of the calibration reticle. Visualization of a 1 Hz square wave between these two levels did not indicate any delay or hysteresis. **C**) The tympanic membrane (TM) is caudo-lateral and ventral to the eye and its overlying skin is naturally pigmented with alternating green and yellow stripes. This lateral view shows a living *Trachemys scripta elegans*, in which two concentric circular skin folds mark the underlying TM under a patch of red pigment. That red stripe has led to this species common name, the red-ear pond slider. **D** and **E** are photographs made through a dissecting microscope directly above a reduced hemi-head preparation, in which the rostral bones in front of the temporal bones (the jaw, palate, orbit and nose) and the rostral telencephalic brain and its adjacent eyes and olfactory nerves. **D** shows the medial side with a recording micropipette whose tip is directed to the brainstem’s cochlear nucleus (red arrowhead) near its attachment to the eighth cranial nerve root from the temporal bone. Brain regions are labeled (S.C, spinal cord; Obex, caudal end of the fourth ventricle; hemi-Cb, cerebellum cut along its sagittal midline and deflected laterally about its peduncle connections to the brainstem; and the midbrain’s Tectum). To stabilize these brain structures, small pins can be seen attaching nearby brainstem tissues to the underlying Sylgard substrate. More rostral parts of the brain are not part of this reduced preparation. **E** shows the lateral side of the preparation, where the sealed tip of a glass probe nearly touched the TM’s central skin. The other end of the glass probe is mounted to a piezo linear actuator, as illustrated in **A**, which is driven electrically based the timing of a voltage pulse to the piezo driver amplifier. **F)** Averaged voltage responses to linear TM movements. In these five traces, small electrical artifacts can be seen when the stimulus pushes the TM in and returns out of the TM (note the arrows below). No response was observed when the sealed pipette tip moved close to the TM without touching it, but small responses were elicited when piezo driven pulses barely deflected the skin covering the TM surface. Large responses to those same pulses occurred when the pipette tip was positioned deeper into the TM surface. **G**) Averaged voltage traces evoked waveforms that were dependent on the piezo driver amplifier’s output polarity and pulse duration (10-ms in this case). Prior to recording, the probe rested deep on the TM. A voltage response occurred when the probe first moved medially and then laterally. Flipping the amplifier polarity elicited an initial lateral movement and then a medial movement. The brainstem response had a different waveform. **H**) The brainstem’s averaged response continued longer than the duration of the electric pulse to the driver amplifier (see also **F** and **G**). Using a constant duration pulse, increasing the gain of the piezo driver amplifier, from low (41) to middle (94) to high (154), resulted in larger and larger linear TM movements and elicited larger voltage responses. The highest gain of the piezo driver amplifier display (peak output voltage of 154 volts of piezo driver amplifier), corresponds to a ∼7 mm linear movement and a large response (lowest trace of **H**).

To prepare this *in vitro* preparation, the cranium was hemisected along the mid-sagittal plane through the base on the cranium to generate a half-head preparation because those responses are best found ipsilateral to the stimulated tympanic membrane. The rostral bones in front of the temporal bones (the jaw, palate, orbit and nose) and the rostral telencephalic brain and its adjacent eyes and olfactory nerves were then removed. The remaining preparation was then hemisected, leaving the brainstem attached to the lateral head, with its labyrinthine sensory organs intact and functional. These tissues were secured, dorsal side up, by a brass strap mounted across the recording chamber to ensure their physical stability during independent movements of the nearby flexible tympanic membrane (Figure 1D, see also Figure 2 A and B). Parts of the midbrain, cerebellum and spinal cord remained connected to facilitate physical stability of the neural tissue by pinning these brainstem structures to the Sylgard-coated floor of the recording chamber (Sylgard-184 Silicone Elastomer, Dow Corning, Midland, MI). Extra care was taken not to grasp the temporal bones by the bilateral bony depressions that are the two tympanic membranes, because they are each connected to their underlying thin columella bones that relay vibrations of each tympanic membrane to each otic capsule. Damage to the delicate columellae and the flexible hydraulic seals of their end-feet onto the oval windows of each otic capsule might damage the relay of TM vibrations, not to mention causing harm to the sensitive stereocilia of the auditory hair cells therein.

**Figure 2.**
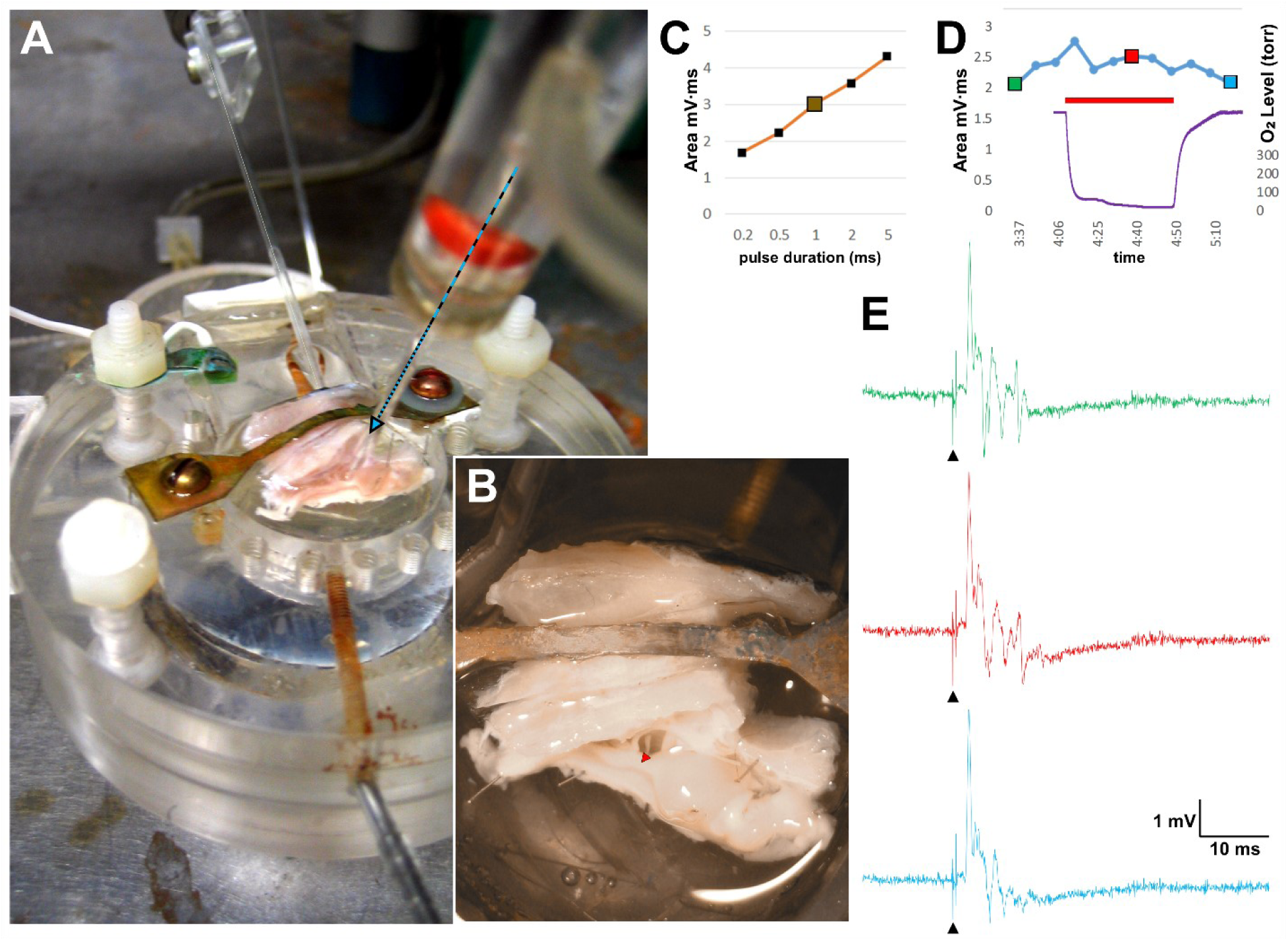
Example of Effect of Hypoxic Media on Responses to TM movement pulses. **A**) Photograph of stimulus and recording set-up (also see Figure-Legend 1**A** and Methods). The recording/perfusion chamber consists of an inlet stainless steel tube in foreground, connected to a nearby stopcock to direct either normoxic or anoxic perfusate into the chamber. **B**) A magnified view of **A** shows the reduced half-head preparation of an *in vitro* left temporal bone and its attached brainstem, secured to the chamber by a brass strap and fine stainless-steel pins, respectively. Seen above the preparation in **A** from left to right is the glass shank of the TM stimulator, the out-of-focus suction pipette removing the perfusate, and the recording micropipette (blue dashed arrow) directed to the lateral brainstem. In **B**, a small dark oval gap shows the space between the inner wall of the temporal bone and the lateral brainstem, within which is the root of the eight cranial nerve. Note the recording site is marked with a red arrowhead pointing down-left. **C**) Plot of the response rectified area (mV*ms) recorded in the brainstem as a function of the duration of the pulse that moves the TM stimulator. **D**) A plot shows time on the abscissa. The red horizontal line indicates the 35 min application of hypoxia, during which the lowest P_O2_ value was 17 torr. The left ordinate axis plots the responses to 0.5-ms TM pulses (blue line) and the right ordinate axis shows the P_O2_ torr values measured within the perfusion chamber (purple line). **E**) Three voltage traces of brainstem responses to 1-ms stimulus pulses (green during control, normoxic media; red during hypoxic media; and blue during recovery; normoxic media). The responses of those traces were plotted using the corresponding colors of the large symbols in **D**.

The tympanum was vibrated with a smooth glass probe mounted on a piezo-electric motor (PZL-007 linear actuator, Burleigh, Fishers, NY). The probe, which was fashioned from a 2-mm glass capillary tube whose tip was sealed by fire-polishing, was placed gently on the central region of the TM so as not to damage the underlying mechanical auditory structures (Figures 1A and E). The tip position of the glass probe was adjusted by a micromanipulator to touch the skin surface covering the TM itself, while visually observed through a high-magnification dissecting microscope (Olympus SZX12 Stereo Microscope, Center Valley, PA). The piezo-electric linear actuator moved the probe up to 7-µm step movements that were roughly orthogonal to the surface of the tympanic membrane as driven by a PZM 150-volt driver amplifier (Burleigh Park, Fishers, NY). The stepper is rated to have a 5% non-linearity, 16% hysteresis and a 4.5 KHz frequency response. A Master-8 Pulse Stimulator (AMPI, Jerusalem Israel) controlled the driver of the piezo actuator.

To calibrate the excursion of the stimulating probe that pushed on the TM, a stiff tungsten rod was fixed to the piezoelectric actuator in place of the almost transparent glass probe and aligned on a reticle on a microscope stage beneath a high-magnification microscope. The linear position of this opaque rod was photographed at the extremes of piezo steps (Figure 1B). The tip of the tungsten rod, analogous to the glass probe, transversed ∼7 microns, as expected from the PZL-007 piezo-electric linear actuator. Preliminary experiments indicated that mechanical steps that pushed onto a correctly positioned TM would evoke a brainstem field potential at fractions of the maximal distance of 7 µm (also see (Christensen-Dalsgaard et al., 2012)). This range of movement is consistent with the dimensions of turtle hair cells of *T.s elegans* whose stereocilia are as tall as 10 μm and are estimated to be deflected roughly 100 to 200 nm (Nam et al., 2019; Spoon and Grant, 2013).

To verify that the field potentials were responding to the TM movements, voltages were recorded in the lateral brainstem at or near the region of the cochlear nuclei. The final placement of the micropipette tip was then positioned by a micromanipulator during piezo steps of the TM every three seconds to identify a location that evoked time-locked, large deflections in the voltage trace (e.g., red arrowhead in Figure 2B). In general, the response onset latency was a few milliseconds and the response amplitude of the averaged field potential increased in a graded manner as the TM displacement amplitude increased or the stimulus duration increased. The size and shape of these evoked averaged field potentials were measured as a function of the movement parameters evoked by the piezo-electric pusher onto the tympanic membrane (see Figure 1 F-H). These brainstem responses to the glass probe steps demonstrated the physiological normal aspects of the brainstem auditory responses. *T.s elegans* are known to respond to sounds within both an aquatic and terrestrial environment, with higher auditory sensitivity relative to turtles that live only aquatic or terrestrial lives (Zeyl and Johnston, 2015). Consistent with hair cell mechanoreception in other species, TM movements in these experiments responded to high frequency stimulation with very short latencies (Figure S1), have waveforms that differ based on TM movement direction (Figure 1G), and have larger response amplitudes for increasing movement amplitude (Figure 1H). At stimulus amplitudes of 7-µm, responses do not diminish with repetition, so appear not to damage the stereocilia.

### Preparation 2: Visual Recordings from the Optic Tectum (OT)

The procedures used for recording the *in vitro* visual responses have been described in several publications, for which neural recordings were made in visual brainstem structures including pretectum, the nuclei of the accessory optic system, the optic tectum, the nucleus isthmi and the cerebellum (Ariel and Fan, 1993; Ariel and Johny, 2007; Fan et al., 1995; Kogo et al., 1998; Rosenberg and Ariel, 1990; Saha et al., 2010). The present study examined only the brainstem responses from the optic tectum. In brief, once the jaw and attached muscles were removed, the eyelids, conjunctivae and bone that formed the orbits were dissected away from each eye. The cranium was opened and the brain + eyes were removed from all the bones. The entire telencephalon was cut from the thalamus and discarded. Eyecups were formed by slicing each eye along its anteroposterior midline, followed by the removal of the lens and excess vitreous humor. The entire preparation of both eyecups attached to the brainstem was submerged in control media in a recording chamber, with both the retinas facing upward, filled with oxygenated media and exposed to the diffuse bright-white light from a light-emitting diode (LED; driven by a 5-volt pulse from the output of Digidata 1320A controlled by ClampEx 9 software, Axon Instruments, Foster City, CA).

These experiments used LED pulse durations as a measure of stimulus light intensity. Stimulus duration was controlled by voltage steps with great accuracy given the temporal nature of LEDs of their nearly instantaneous onset and offset of light pulses. Phototransduction, on the other hand, proceeds very slowly, especially for poikilothermic animals like aquatic turtles spending their winters in near-freezing environments. Therefore, turtle photoreceptors integrate individual photon absorptions as transduction events within a long time window of hundreds of milliseconds for cold conditions, especially for rod photoreceptors. At warmer temperatures, integration times may be as short as 100 ms for cone photoreceptors (Baylor and Hodgkin, 1973) (Granda et al., 1986).

For recording of the field potential responses in the brainstem, the OT of the dorsal midbrain was placed upward in the recording chamber, and its overlying meninges were carefully peeled away to permit a recording micropipette tip to smoothly penetrate the tectal surface to a depth of approximately one-half millimeter. The field potentials recorded there represent the aggregate, low-pass filtered spike activity near the electrode tip in the tectum (Figure S4).

In some cases, the surgery first created the eye + brainstem preparation, while preserving the caudal head tissue for use in a separate preparation that included the tympanic membranes and the temporal bones as described above. The various preparations were secured into plexiglass chambers containing a continuous flow of oxygenated perfusate. The brainstem was affixed by several pins to the Sylgard floor of the recording chambers (see details in Rosenberg and Ariel 1990). During each recording session, a wide range of stimulus intensities were presented to determine the range of intensities (0.05 – 10 ms) that elicited electrical field responses that were consistently above response threshold and below response saturation. Light intensities within that range (1 – 5 ms) were selected to monitor the visual responses during the normoxic and severe hypoxic aCSF conditions.

Prior to the onset of severe hypoxia, control responses were recorded to establish a stable baseline for the sensory responses under oxygenated conditions. Then, a stopcock located close to the recording chamber switched the perfusate from the oxygenated media (95% O_2_/5% CO_2_) to severely hypoxic media (95% N_2_/% 5% CO_2_). To minimize contamination of the perfusate with atmospheric oxygen, the tubing carrying the media from the reservoir to the recording chamber had low oxygen-permeability (Versilon™-c-210 tubing; Oxygen Coefficient 18 • 10^-11^ • cc • cm / cm2 • s • cmHg; Saint-Gobain, Akron, OH) or PTFE tubing ensheathed in an outer tube filled with flowing N_2_ gas. In this OT preparation, as with preparation 1 for auditory recordings, the recording chamber was covered with a thin cover-glass to block air diffusion into the chamber’s perfusate. Also, with both brainstem preparations, recording from either the cochlear nuclei or the OT, the overlying manipulators and thin cover-glass were removed after each experiment to permit unobstructed overhead photography of the tissues within their recording chambers. The images of those reduced preparations were clearly observed either submerged with the perfusate or immediately after drainage of the perfusion media.

### Preparation 3: Visual Recordings from the Retinal Eyecup

To isolate the tectal responses from those of the retina itself, we determined the effects of severe hypoxia on the electroretinograms (ERGs) from single eyes that were separated from the brainstem by severing them from their optic nerves. In contrast to the brainstem recordings described in the two preparations above, which produce relatively small voltage responses, visually evoked ERGs produce large responses measured in an intact eye where the corneal/vitreal side of the retina is electrically isolated from the scleral side of the retina. However, in these ERG experiments, the eye was hemisected to form an eyecup (Figure S2B) for application of anoxic aCSF, creating a potential conductive path that could shunt the radial retinal current between the two sides of the retina if submerged. This shunting path was minimized within the special eyecup recording chamber and the cap for that chamber as follows..

In vitro ERG recordings require that the eyecup be placed in a gaseous environment so that the field potential between the retina’s inner surface is electrically isolated from its scleral surface. To this end, a special eye chamber was fabricated to support the eyecup vitreal side up in a humidified gaseous environment. The chamber as covered by a cap (shown in the foreground of Figure S2A) through which enters the perfusion tubing, the tips of a 1.5 mm glass recording micropipette and a 3 mm O2-optode (PyroScience GmbH from Ohio Lumex Co) into the vitreous of the eyecup. While the voltage was measured between the micropipette tip in the vitreous with respect to a ground wire touching the scleral side of the retina, ERGs were evoked by light flashes from the overlying Light-Emitting-Diode. In this configuration, the recorded in vitro ERG of the intact eyecup had all the characteristics of an ERG of the in vivo eye.

Maintaining hypoxia in the special eye chamber was difficult due to the several small holes in the transparent plastic cap necessary for access of electrodes and perfusion tubing to enter the chamber. Those access holes provided limited and unwanted exposure to ambient air. Therefore, an additional small hole was made in the eye chamber’s cap to permit a thin needle to deliver a continuously gas flow of 95%-N2/5%-CO_2_ to flush out the oxygen contamination from the ambient air. This approach to maintain hypoxia in the retina and limit oxygen contamination was successful to a great extent. ERGs were recorded while the O_2_-optode recorded very low oxygen torr. However, in some cases over time, air would enter the eyecup chamber and reach the retinal eyecup via the media bathing the vitreal surface, causing oxygen contamination and slow drifts of increasing P_O2_. Gradual increases in P_O2_ may also be due to the high metabolic activity of the retina due to its own respiration (Özugur et al., 2020). Note that these P_O2_ positive drifts up to 150 torr were not observed in the experiments for preparations 1 or 3, when the brainstem chamber was totally filled with physiological perfusate and covered with a thin glass plate.

### Electrophysiological Techniques

All recording micropipettes were pulled from 1.5 mm OD thick-walled borosilicate glass using a horizontal puller (P-87; Sutter Instruments, Novato, CA) and filled with 3 M potassium acetate, resulting in 0.5-1.8 mΩ microelectrodes. The amplified signals were recorded in the current clamp mode of an Axoclamp 2B amplifier and further pre-amplified (Warner Instruments IPF-200; c/o Harvard Bioscience, Holliston, MA) prior to digitization (ClampEx using Digidata). Sensory stimuli (single pulses < 20 ms or brief trains of pulses) were recorded either as single traces or averages of up to 64 traces. At least 3 seconds separated each stimulus. The ClampEx software sampled those voltages with a 50-µs sampling interval, averaged many dozens of these individual stimuli or pulse trains and then saved those traces as digital files. All digitized traces generated were analyzed offline using Clampfit (Axon Instruments, Foster City, CA). The waveforms of the field potential responses varied with the micropipette position in the brainstem and were multi-phasic in nature. Moreover, by using an amplifier with only a high-frequency cut-off filter (50 KHz), the baseline value of each trace differed slightly from trace to trace. Therefore, the average value of pre-stimulus component of each voltage trace was digitally subtracted and the post-stimulus waveform was rectified at the computed baseline value. Using these filtered traces, amplitudes were quantified as the areas under the curve in mV·ms for the recordings made from the cochlear nuclei and the OT.

The analysis of ERG responses differed from those recorded in the brainstem because of the two consistent ERG components: an a-wave (hyperpolarization) and b-wave (depolarization) (Figure S3B). Six different stimulus intensities were presented in an increasing order of brightness, from near-threshold stimuli to just-saturating stimuli. The ERG traces were quantified using a standard set of formal waveform parameters using the Clampfit software: latency to a-wave onset, latency to a-wave peak, a-wave peak amplitude, a-wave area, b-wave peak amplitude, b-wave area, time between a-wave and b-wave peaks, latency to b-wave peak, maximum slope between a-wave and b-wave, and the time of maximum slope (Figure S3A).

### Statistical Analysis

Single factor ANOVA was used to determine if severe hypoxia and reoxygenation affected the evoked responses of the cochlear nuclei and the OT. For the ERGS, two-factor ANOVA was used to elucidate the effects of both stimulus intensity and oxygenation state, and to determine if an interaction between the variables existed. Whenever equal variance or normality assumptions were violated, the data were either transformed or ranked prior to testing. Statistically significant single factor effects and interactions were further elucidated post-hoc with Student-Newman Keuls testing. All analyses were conducted using Sigmaplot 13 (Systat Software Inc., Chicago, IL). The significance level was always alpha < 0.05.

## Results

In the present study, the auditory and visual responses in a reduced brain preparation were examined to evaluate whether the effects of hypoxia on sensory systems would differ depending on the mechanisms that transduce the two environmental stimuli into neural messages within the brain. The first set of experiments studied the mechanoreceptive hair cells of the auditory system and the second set studied the light-sensitive photoreceptors of the visual system.

### Brainstem Responses to TM movement steps are unaffected by bathing in near-anoxic media

To determine whether near anoxic superfusion affects the mechanosensory functions of the turtle inner ear, voltage recordings were made from the brainstem with a low impedance (1-3 MΩ) glass micropipette placed near its connection with the eighth cranial nerve during mechanical pulses of TM (see Methods, *preparation 1* and Figure 1). A representative response of the evoked potential to near anoxic superfusion is depicted in Figure 2 and mean responses from eight experiments in Figure 3. The Po_2_ of the superfusate decreased to a 10.1 ± 1.5 torr (mean ±SEM; range 2.9 - 20.3). Despite this, there were no statistically detectable effects of prolonged near anoxia on the responses to TM movements, either for the absolute (Fig. 3B) or normalized (Fig. 3C) response areas. To confirm that we were able to detect a diminished response if it had occurred by another mechanism, we applied lidocaine to the preparation, which strongly attenuated the evoked potential of the TM movements (Fig. 4).

**Figure 3.**
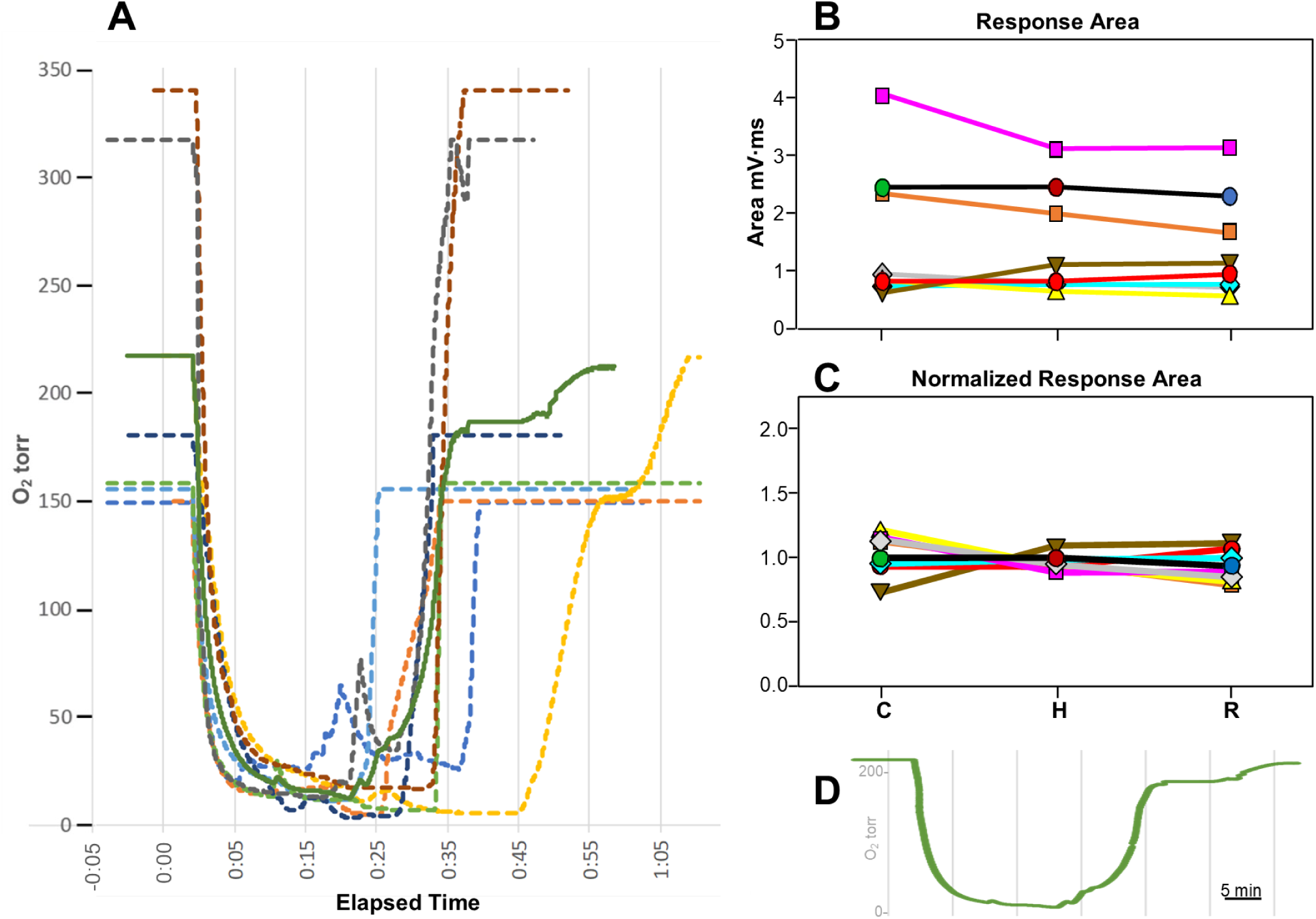
Analysis of Responses to TM movement in Recording of Eight Brainstems during Hypoxic Media measured by Optode Average. In eight experiments, single pulses were delivered to the TM by a piezomotor while recording from the ipsilateral cerebellar peduncle close to the brainstem’s junction with the eighth cranial nerve. **Panel A)** Twenty P_O2_ levels were recorded per second and plotted as dashed lines for each of eight experiments. The rotation of a stopcock directed anoxic perfusate into the recording chamber, resulting in rapid descent of the optode output from the high P_O2_ -bubbled perfusate to the zero-O_2_ perfusate. This plot shows the eight dashed lines that were aligned with the rapid descent of P_O2_ levels. The onsets and durations of the decreases in P_O2_ levels were variable due to differences in perfusion rates, the positions of the hemibrain in the chamber and the experimenter’s time to switch back to the control perfusate. For two of the eight experiments, their dashed lines were shifted because of a slower initial descent of their P_O2_ levels relative to the other six experiments. The re-ascent of the P_O2_ levels depended primarily on the rotation of a stopcock duration by the experimenter back to the normoxic perfusion. The solid green line represents the average of the eight dashed lines, determining three experimental epochs: a control high-O_2_ condition, a hypoxic condition and a condition following return to high-O_2_ after the cessation of N_2_ bubbling. **Panel B** and **C** have two response plots following the TM movement, displayed as response area and normalized response area, respectively. The data points are aligned with the averaged optode levels (shown below in **Panel D,** that were replotted from the green line of **Panel A)** to indicate the sequence of the three conditions of media oxygenation: normoxia, N_2_ bubbling and recovery. Note that the optode data of Figure 2D is shown here in **Panel A** as the dashed brown line. Also, the data points connected by the black lines of the Panels B and C are the same data as shown in the green, red and blue response traces in Figure 2E.

**Figure 4.**
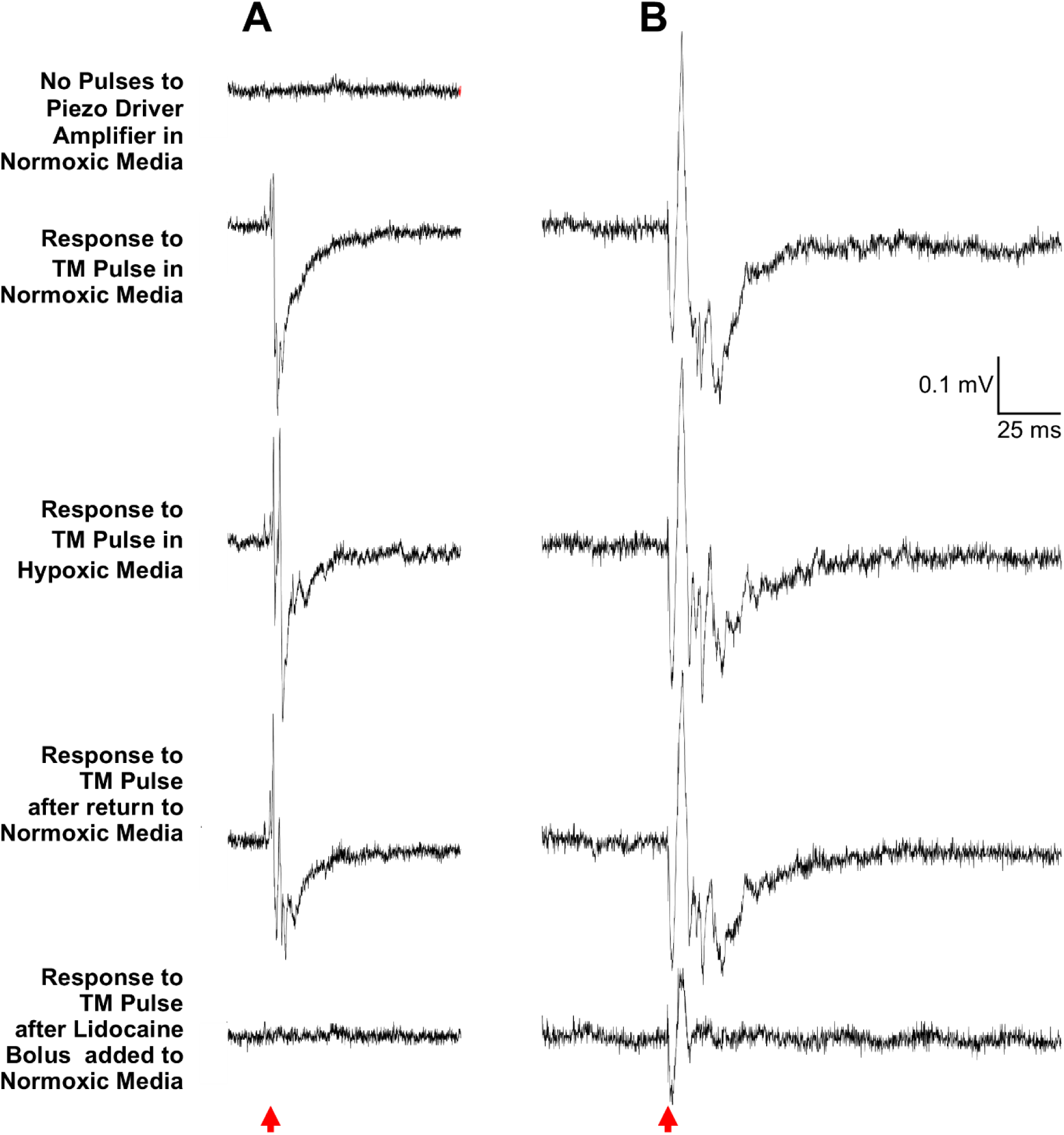
Two examples showing that Voltage Deflections required TM movement and Biological Responses. **A)** In this example, the recorded trace was generated when 64 0.5-ms pulses triggered the voltage recording of 64 traces, yet those pulses were not triggering the piezomotor driver amplifier. The resulting averaged voltage trace in the brainstem did not exhibit synchronized voltage deflections, demonstrating that the field potential responses to TM movements are not artifacts due to the amplifier electronics. When the piezomotor amplifier was being activated by the pulses (indicated below the traces by a red arrowhead), the next three traces showed evoked field potentials that were synchronized to TM movements, whether the neural tissue was bathed in oxygenated or hypoxic media. After the brainstem was again bathed in oxygenated media, a 50-µl bolus of a fresh stock of 40mM lidocaine was added to the estimated 1 mL volume of chamber’s brainstem and perfusate (∼20-fold dilution, leading to an estimated 2 mM final concentration of lidocaine in the chamber). Within a minute of the addition of the lidocaine, the evoked field potentials evoked by TM movements were blocked (see lowest trace). This lower trace is very similar to the top trace, recorded in the absence of TM movement, supporting the conclusion that the field potentials require both a vibration of the middle ear and the biological mechanism to convert that vibration into a neural signal that can reach the brainstem. **B)** In a second example, a 2-ms pulse to piezomotor that vibrates the TM also evoked synchronized field potentials in the brainstem, whether the neural tissue was bathed in oxygenated media, hypoxic media or returned to oxygenated media. However, after a lidocaine bolus was then added to the oxygenated media in the recoding chamber, the neural block that resulted was strong but incomplete. Note that the initial components of the field potential have the same onset latency but that the delayed response components are absent. The partial effect of lidocaine in this example suggests that mechano-transduction within the middle ear and otic capsule is still functioning but that the remaining sensory path within the brain was suppressed by the limited action of lidocaine.

### Electroretinographic (ERG) responses to light flashes are affected by hypoxic media

Due to the surprising lack of an effect of hypoxia on the field potential responses of brainstem neurons produced by from mechano-sensitive hair cells in the turtle’s inner ear, we performed a set of complementary experiments on the visual system, whose input also originates outside the cranium. In the first set, ERGs were used to measure the retina’s visual response as the modulation of the electrical potential at the cornea relative to the sclera. To this end, ERG traces were recorded in response to very brief light flashes from the light-emitting diode of varying durations (0.05, 0.2, 0.5, 2.5, 5.0, 10.0 ms) during perfusion with normoxic media, then during hypoxic media, and finally, after the return to the normoxic media (recovery).

Representative responses of the evoked responses of the retina to the light flashes during hypoxic superfusion are depicted in Figure 5. The Po_2_ of the perfusate decreased to 28.6 and 12.1 torr (Fig. 5A and 5B, respectively), producing modest decreases in the amplitude of the evoked responses with both 0.5 ms (Fig. 5D and 5E) and 5 ms (Fig. 5F and 5G) stimuli. There was an overall effect of hypoxia (single factor effect), independent of stimulus intensity on the peak amplitudes of both a- and b-waves. Similar analyses were performed on data from experiments in which Po_2_ only decreased below 60 torr but not below 30 torr. There too was a single factor effect of moderately hypoxia on ERG parameters of the a- and b-waves (but not b-wave area), suggesting that visual responses may also be decreased even when a turtle may be making a a prolonged dive. We found no statistical interaction between treatment and stimulus intensity, which means the hypoxic suppression was independent of stimulus intensity.

**Figure 5.**
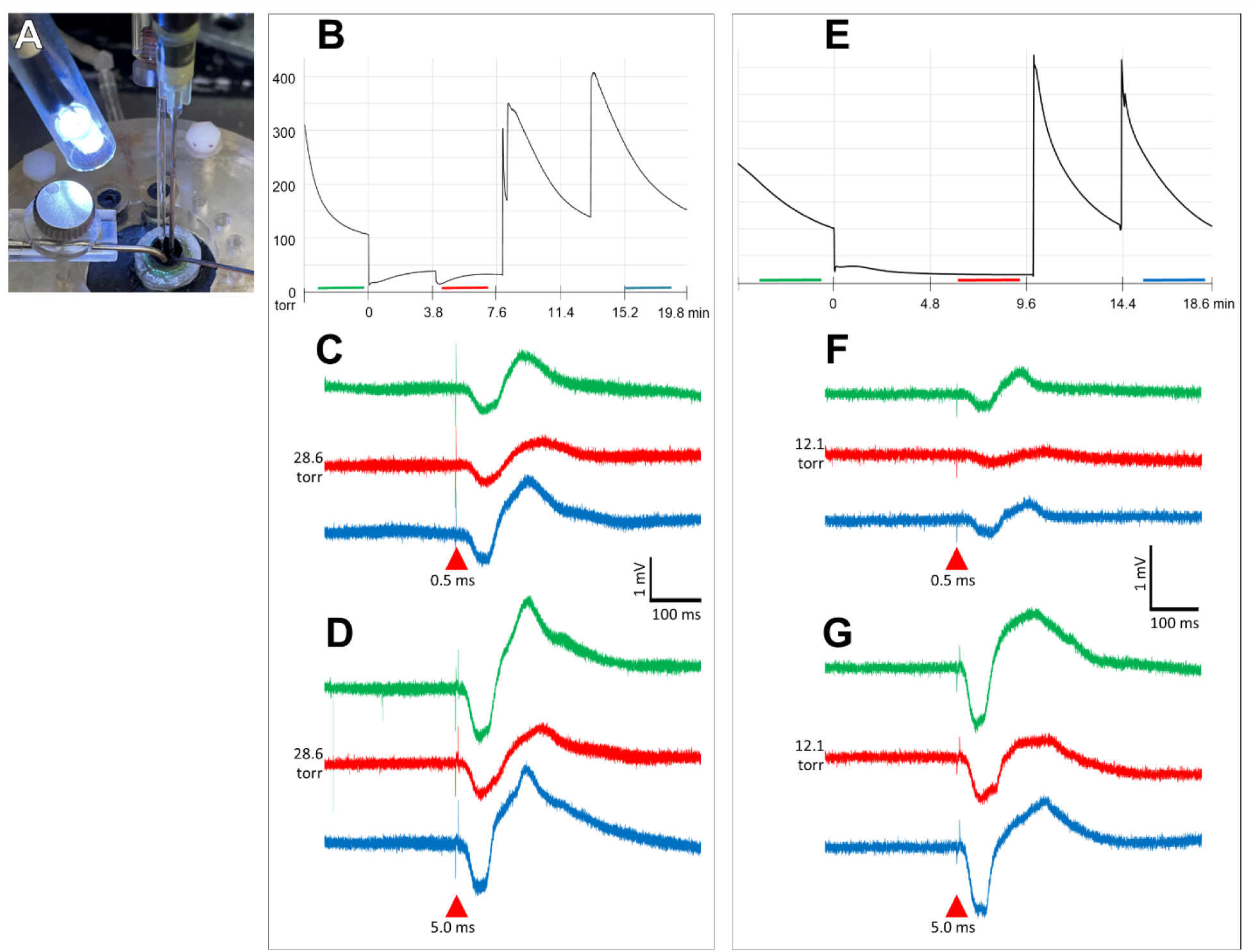
Two Examples of Effect of Hypoxic Media on Responses to Single Light Pulses recorded in the Retinal Eyecup. **A)** Photograph of ERG setup, with micropipette and optode in the eyecup, media application from left curved needle and gas flow from right curved needle. For clarity, chamber cap that was covering eyecup was removed for this photograph. **B**) optode trace of the left example, with horizontal lines showing the times of control, hypoxic and recovery recordings. The red bar under the optode trace indicates that the eyecup media during hypoxia had a P_O2_ of 28.6 torr. **C**) ERG responses of the left example to a low (0.05-ms) stimulus intensity, during control, hypoxic and recovery conditions (traces coded by color). **D**) Similar to **C**, ERG responses of the left example to brighter (5.0-ms) stimulus intensity. Like panels **B** – **D**, the panels **E - G** show similar data for the second example in the right column but, in this second experiment, the hypoxia media in the eyecup dropped down to a P_O2_ of 12.1 torr (**E**, red line). **F** shows ERG responses of this right example to low stimulus intensity (0.05-ms), during control, hypoxic and recovery conditions while **G** shows ERG responses to a brighter (5.0-ms) stimulus intensity.

The quantification of a- and b-wave amplitudes are plotted on the y-axis for the six different stimulus intensities are seen in Figure 6A and 6B, respectively. Those measures are also compared for the conditions of normoxia, hypoxia below 30 torr P_O2_ followed by recovery (C, H, R on the x-axis). As expected, the brighter stimuli evoked larger responses, but for each intensity tested, those responses were decreased during hypoxia.

**Figure 6.**
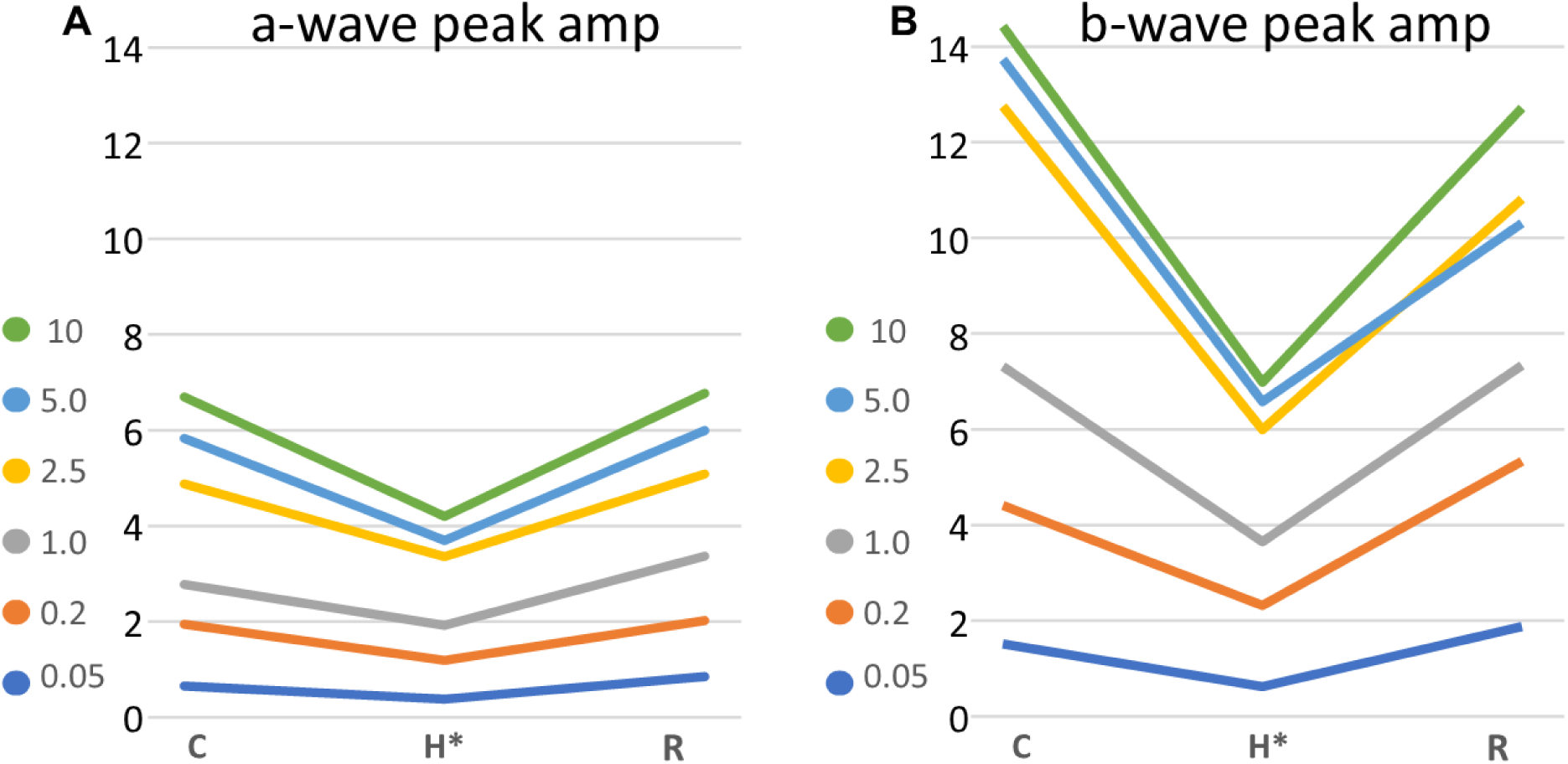
Effect of Hypoxic Media on the ERG Peak Amplitude for each Stimulus Intensity. The ERGs were evoked by six light flash intensities during conditions of normoxia, hypoxia below 30 torr P_O2_ followed by recovery. **A & B** show graphs of the peak amplitudes of a-waves and b-waves, respectively, during control media (**C**), then hypoxic media (**H**, <30 torr pO_2_) and then during recovery (**R**) back to the control media (**C, H, R** on the x-axis). Each line is the average of the response of a single light intensity, averaged from all the seven retinal eyecups. Note that the increasing stimulus strengths resulted in increased responses, but that the responses to all six light intensities were decreased by hypoxia. Also note that the b-wave peak responses are roughly twice as large as that of the a-wave responses.

As the light responses of each eyecup preparation differed due to animal and surgical variability, the quantified data used in Figure 6 are re-sorted into individual experiments and plotted in Figure 7. Figure panels 7A-7D depict the changes in normalized amplitudes of a- and b- wave responses and figure panels 7E-7F depict their timing changes (their onset and implicit times of eyecups from all seven preparations). The values were averaged across stimulus intensities because there was no significant interaction between treatment and stimulus duration. Hypoxic superfusion produced no significant change in the amplitude, onset, and implicit times of the a-wave, but the b-wave showed significant decreases in amplitude (Figs. 7C and 7D) and increased implicit time (Fig. 7 F).

**Figure 7.**
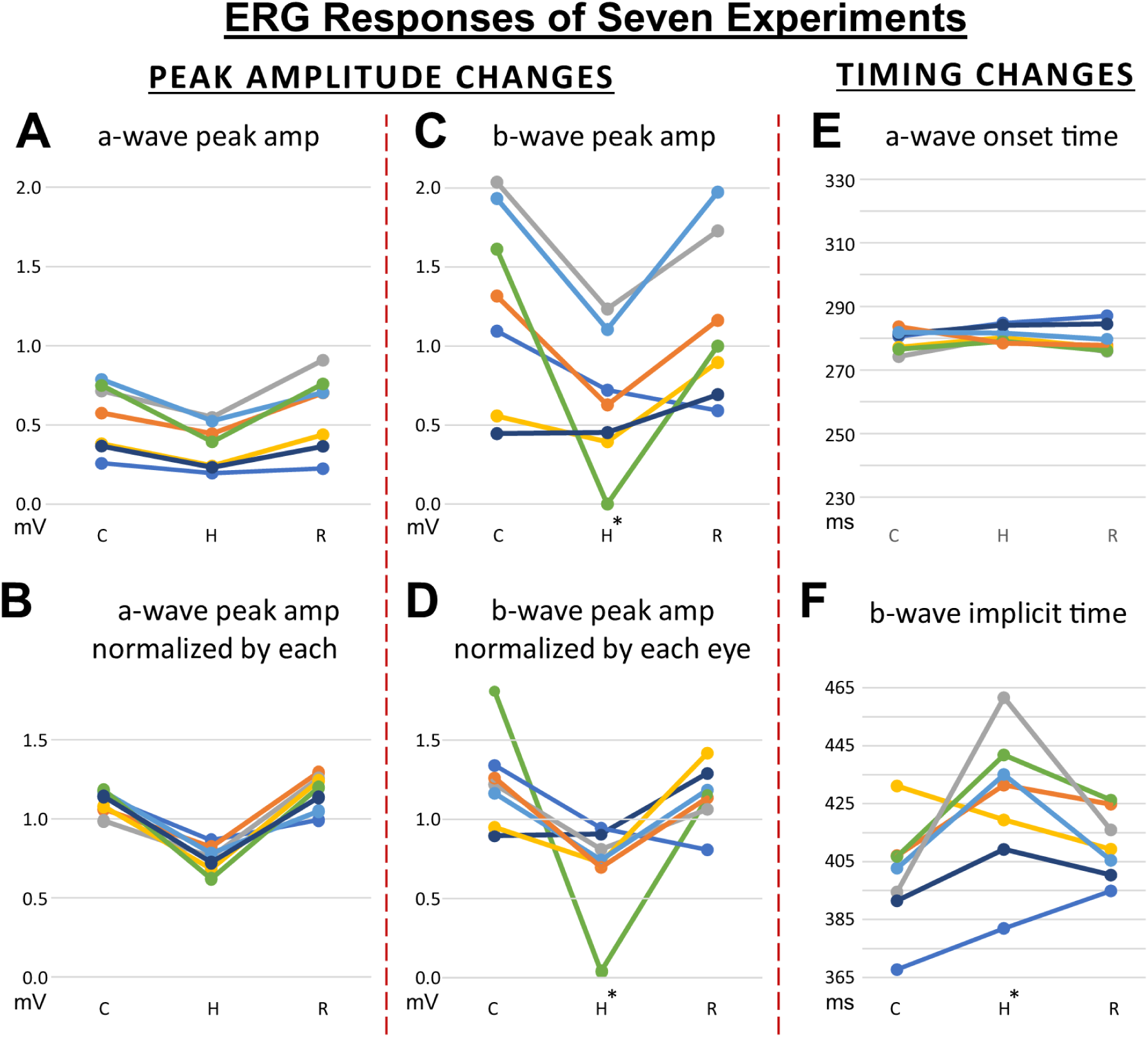
Effect of Hypoxic Media on Peak Amplitudes and Latencies of each of the Seven ERG experiments. A statistical analysis was performed on the ERG peak amplitudes, each colored line showing the result of one of retinal eyecup studied. In the left column, panel **A** shows mean a-wave peak amplitudes for the seven experiments, plotting their responses in millivolts. Below it, Panel **B** displays the same a-wave data but normalized by the all the response of that retinal eyecup. Normalization better displays the similarities of effect of hypoxia on each eyecup studied. In the middle column, panels **C & D** show the same format as the left column but using the b-wave responses. Note that, by comparing the left and middle columns at the same scaling, b-wave responses are larger and more variable for the same seven experimental tissues. Also note that the responses of all seven eyes were decreased by hypoxia (*: p < 0.05). In the right column, timing changes due to hypoxia were analyzed. The data on panels **E & F** reveal that the a-wave timing was not affected by hypoxia, while the implicit times of the b-waves in nearly every experiment were delayed in the presence of hypoxia (*: p < 0.05). During hypoxia, no other b-wave timing measures were affected during hypoxia.

### Recording of OT field potentials respond to retinal light flashes

The second set of visual response experiments involved the recording of brainstem light responses in the OT that originate in the photosensitive retina while superfusing the preparation with hypoxic media. Like the averaged field potentials recorded in the auditory brainstem, averaged field potentials were measured as the area under the rectified curve.

An example of a recording of the light-evoked responses of the OT with hypoxic superfusate is depicted in Figure 8. A moderate amplitude response to a dim stimulus (0.5 ms, tan box in Fig. 8C) was chosen to monitor the responses in different media. As described above for the graph in figure 2D, the blue line of Figure 8D shows the quantified responses to evoked rectified field potentials. Examples of raw traces from this experiment are shown in Figure 8E (control normoxia, green; hypoxia, red, recovery, blue), which correspond to the colored symbols in of Figure 8D. Within minutes of starting the anoxic perfusion, the Po_2_ of the superfusate decreased to less than 10 torr, maximally suppressing the evoked response of the OT within ∼10 minutes. Upon returning to normoxic media, both light responses and Po_2_ returned to control levels, also within ∼10 minutes.

**Figure 8.**
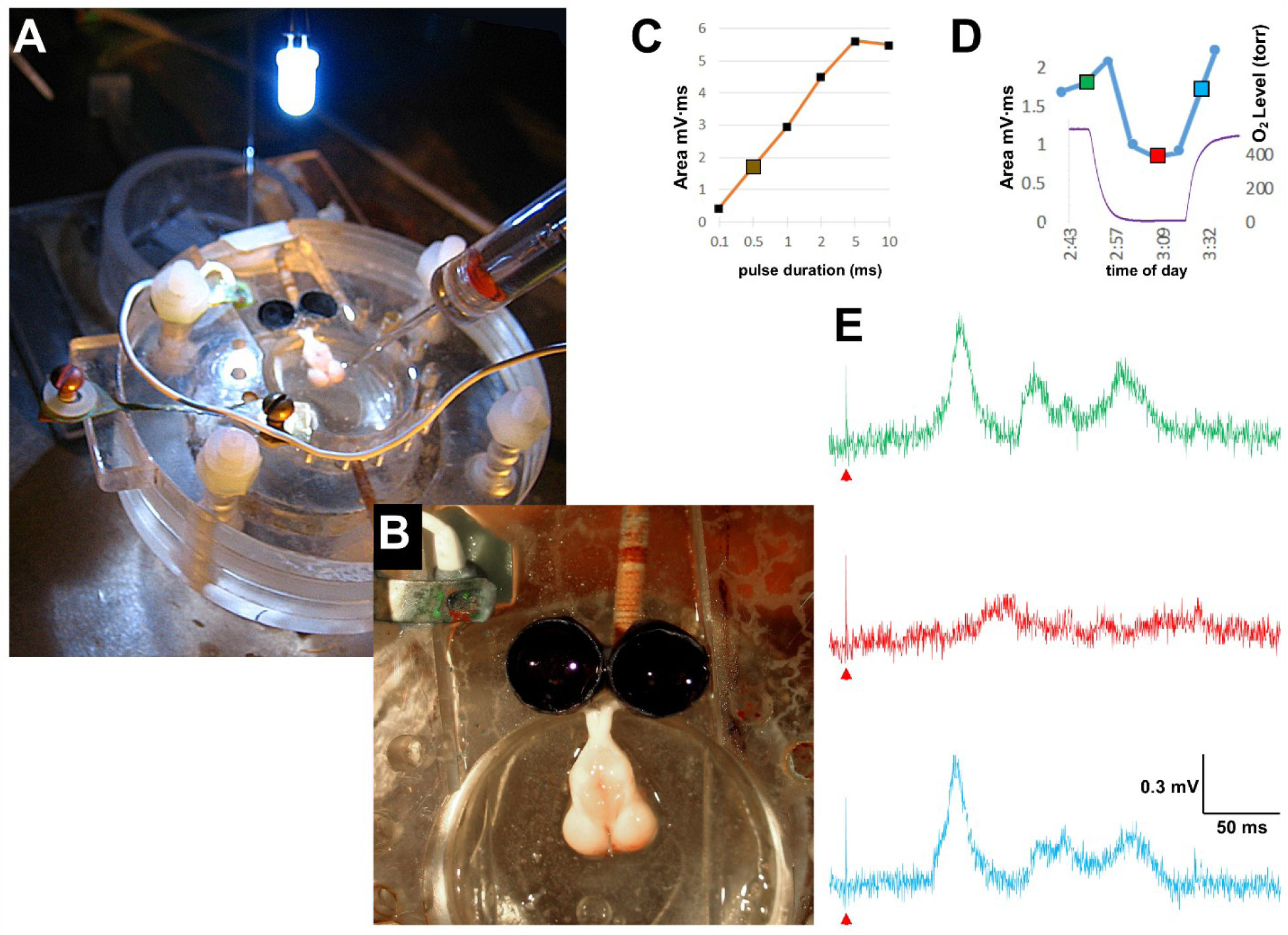
Responses to Full-Field Light Pulses to the Retina: OT Recording during Brainstem Hypoxia. **A**) Photograph of the same recording/perfusion chamber as described in figure 3, but this experiment studied responses of OT to 0.5-ms LED flashes above the two eyecups, both connected to the midbrain via the optic chiasm. The recording micropipette was placed just below the dorsal surface of the right tectum. A stopcock directed either normoxic or anoxic perfusate into the chamber. **B**) A magnified view of the reduced preparation seen from above. **C**) Plot of the response amplitudes recorded in the tectum as a function of the stimulus brightness. **D**) A plot shows time on the abscissa (with the red line indicating the 26 min application of hypoxia). The left ordinate axis plots the amplitude of tectal responses to 1-ms pulses (blue line) and the right ordinate axis shows the P_O2_ torr values measured within the perfusion chamber (purple line). Seven torr was lowest value of P_O2_ during the perfusion of hypoxic media. See Text for more details.). **E**) Three voltage traces of tectal responses to 1-ms stimulus pulses (green, control responses in normoxic media; red, during hypoxia media bubbled with N_2_; blue, recovery after return to normoxic media). These voltage traces show responses to the stimulus intensity that corresponds to enlarged symbol in panel **C**, as well as the large data point of the blue line of panel **D**.

Like Figure 3, Panel A of Figure 9 depicts overlapping dashed lines of optode traces of four of the five OT experiments. In Panels B and C, the absolute and normalized response area values from all five preparations are plotted, respectively, below which are the averaged optode plots (derived from the solid green line of Panel A). During hypoxic superfusion, the responses significantly decreased to ∼75% of control levels, and then fully recovered upon reoxygenation (*: p < 0.05).

**Figure 9.**
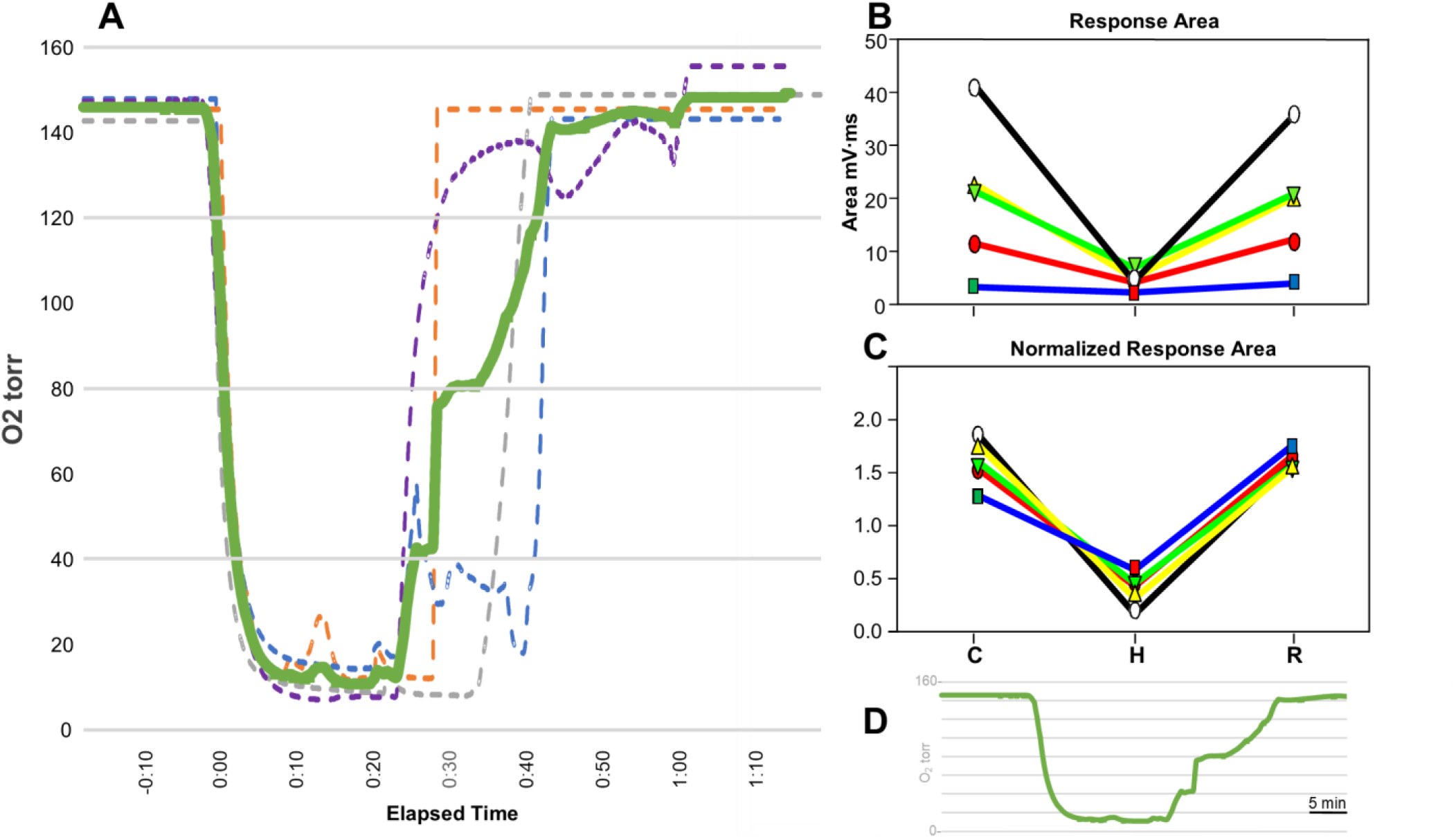
Statistical Analysis of Five OT Responses to Retinal Light Flashes during Brainstem Hypoxia. **Panel A)** Similar to figure 3, four P_O2_ levels were recorded per second and plotted as dashed lines. These dashed lines were aligned with their rapid descent following the switch from the high O_2_-bubbled perfusate to the zero-O_2_ media into the recording chamber. The re-ascent of the O_2_ levels varied depended on the time that normoxia was reapplied by the experimenter. The solid green line (also shown in **D**, represents the average of the four dashed lines, and determines three experimental epochs: a control high-O_2_ condition, a hypoxic condition and a condition following return to high-O_2_ after the cessation of N_2_ bubbling. **Panel B** and **C** have two response plots due to the light flashes, displayed as response area and normalized response area, respectively. The data points are presented over the averaged optode levels (**Panel D)** to indicate the sequence of the three conditions of media oxygenation: control, hypoxic (N_2_ bubbling) and recovery. The optode data of figure 8E is shown in **Panel A** as the dashed gray line. Also, the colored traces in Figure 8D are displayed as the response amplitude data points in **Panel B** and **C** filled with the same colors: green, red, and blue. Note that there was a clear reduction in the OT’s light responses during the strong hypoxia, unlike the response amplitudes during a similar application of strong hypoxia when recording the responses to TM movements at the brainstem’s junction with the eighth cranial nerve, as shown in figure 4.

Unlike the short onset latency of brainstem responses of a few milliseconds following TM movement (see Figure S1), the OT light responses exhibited much slower onset latencies of several dozen milliseconds (Figure S4-S6). It is therefore not surprising that the brainstem responses to trains will respond to individual TM stimuli of interstimulus intervals as short as 2-ms (see also Christensen-Dalsgaard et al., 2012), whereas OT responses to individual flashes within trains blend together, exhibiting temporal response summation even when intervals are as large as 200-ms (Figure S6).

The onset latencies of OT primary depolarizing wave were also significantly longer than ERG responses, averaging 50 ms for moderate light intensities (Figure S5A), compared to 14 and 45 ms for the a-wave and b-wave, respectively. Like the ERGs, the onset latency of the OT responses decreased as the light stimulus intensity decreased, to as long as 105 ms (see Figure S5). In contrast, brighter light flashes elicited a second depolarization that grew larger for increasing stimulus intensities. Finally, as shown in Figure 7F, the peak latencies of ERG b-waves slowed during hypoxia, a similar observation was made slowed latencies of OT responses during hypoxia (Figure 8E). These effects of timing may not reflect the effects of hypoxia on sensory transduction mechanisms found in the anoxic-tolerant turtle, in that the onset and peak timing of responses to TM pulses and ERG a-waves were not affected by hypoxia in these data presented above. More likely, these are features of higher brain processing.

## DISCUSSION

This study explored the effects of oxygen deprivation on the sensory brainstem responses recorded in pond turtles, one of the most anoxia-tolerant groups of vertebrates known to science. Utilizing an *in vitro* brain preparation, we demonstrated that the auditory and visual systems exhibit quite different responses to hypoxic and near-anoxic conditions. For the auditory system, responses could still be elicited by controlled, periodic movements of the tympanum even when oxygen tension of the perfusion medium fell below 7 torr. In contrast, but not surprisingly, the visual system showed a high sensitivity to reduced oxygen in the perfusate for both retina and especially OT. Indeed, even relatively moderate oxygen tensions (∼30 torr) diminished the ERG responses. We conclude that the turtle’s visual system has little to no function under anoxic conditions, and perhaps even during some extended voluntary dives. In contrast, at least at the mechano-transduction level, i.e., the cochlear hair cells, auditory function was maintained.

### Brainstem Responses to Vibration of the Tympanic Membrane are Insensitive to Severe Hypoxia

Field potentials responses to TM movements persisted during anoxia. Is this a common finding for auditory brainstem responses in all animals? Similar mammalian field potentials are recorded in vivo, known as the ABR, Auditory Brainstem Response. The ABR is measured in humans clinically and in mammals in laboratory settings (Attias et al., 1990; Skoe and Tufts, 2018). It is a far-field recording of very small neural events, recorded with low impedance surface electrodes on the skin. The averaged waveforms have identifiable morphology, divided into waves I–VI which reflect different structures such as the olivo-cochlear bundle (wave I), the cochlear nucleus (wave II) and the inferior colliculus (wave V) (Kileny et al., 2021). Unlike ABR, the turtle electrophysiological measure of auditory responses used an in vitro technique that is distinct from that of published reports using mammals. The turtle responses were evoked by direct vibration of the TM and were recorded using microelectrodes that had much smaller recording pipette tips of high electrical resistances. Moreover, the effect of anoxia in turtles used a direct brain perfusion and a local measurement in that perfusion media by an optode. In contrast, the mammalian ABR recordings relied on a systemic means to lower oxygen and an indirect measure to compute those oxygen levels (Attias et al., 1990; Haupt et al., 1993; Sawada et al., 2001; Sohmer and Freeman, 2001).

Using ABR, hypoxia created by ventilating the lungs with 6-8% oxygen has been reported to cause a reversible hearing loss in several mammals (Lawrence et al., 1975; Sohmer et al., 1989,1986a,1986b; Hildesheimer et al., 1987; Sohmer and Freeman, 1991; Gycowicz et al., 1988). In guinea pigs, perilymphatic P_O2_ was directly measure during this systemic hypoxia and showed a ∼70% drop with a concurrent decrease of ∼79% of the ABR. Both measures returned to normal during reventilation with room air (Haupt et al., 1993). Using the same animal model yet applying mild hypoxia for a longer duration, it was observed that inner hair-cells were first affected, followed by outer hair-cells, suggesting that inner hair-cells are more vulnerable to hypoxia (Sawada et al., 2001; Shirane and Harrison, 1987). A more recent report measured wave I of the ABR in anesthetized, paralyzed cats, and concluded that hypoxia caused ionic changes within the endolymph, which reduced mechano-transduction by the *Inner Hair Cells* and depressed the strength of the cochlear amplifier of the *Outer Hair Cells* (Freeman et al., 1995).

Similar ABR analyses were performed in fetal mammals, including humans, using more physiological conditions of hypoxia; in this case oxygen delivered by placental diffusion instead of by pulmonary oxygen diffusion following birth. By measuring the average ABR thresholds to bone conducted sounds in near full-term guinea pigs, their neonatal threshold was better than that in the full-term fetus by 20 dB for the same animal (Sohmer et al., 1994b). For the near full-term human fetus, the threshold of its response to vibro-acoustic stimulation was lower and/or that response was stronger when the mother was breathing oxygen relative to breathing room air. At birth, the auditory threshold improved when the new-born breathed air directly (Freeman et al., 1995; Sohmer and Freeman, 2001; Sohmer et al., 1994a). In summary, hypoxia effects inner hair cells more than outer hair cells. Therefore, mammalian brainstem auditory responses are sensitive to weak levels of hypoxia, even those found in the physiological range just prior to delivery. Similarly, latencies of human auditory responses measured more centrally in the brain are known to be increased during weak hypoxia (Fowler and Lindeis, 1992).

Hypoxia also depressed auditory responses in non-mammals, even the crucian carp that is anoxia-resistant (Ping and Jenkins, 1978; Weber, 2016)). Hypoxia was simply produced by stopping water flowing over the gills for up to 20 minutes. During that hypoxia, auditory responses of spiking axons in the auditory nerve would decrease or be totally blocked (see figures 5 & 6 of Fay and Ream, 1992).

### Auditory responses of turtles on land or submerged in a pond

Although pond turtles are uniquely adapted to both terrestrial and aquatic life, hypoxic environments are predominant in eutrophic habitats like muddy bogs where pond turtles frequently live. In an experiment that studied auditory thresholds of turtles, either submerged or airborne, head withdrawal behaviors were measured to different intensities of sounds paired with an aversive stimulus (Patterson, 1966). This behavioral response showed that turtles were more sensitive to underwater stimuli than to airborne stimuli. Similarly, ABR was measured in airborne and underwater turtles and their sound thresholds in water were 20-30 dB lower than those measured for airborne sounds (Christensen-Dalsgaard et al., 2012). Taken together, a pond turtle’s hearing is very likely to be effective while hypoxic underwater due to increase auditory sensitivity in that environment.

It is important that we point that our findings are limited to more peripheral stages of the auditory system, where: 1) the otic capsule and temporal bone were both bone ischemic, eliminating possible humoral influences, 2) that the production of healthy endolymph necessary for hair cell mechanotransduction was functional even when bathed in hypoxic media and 3) that hypoxic hair cells maintained synaptic communication to ganglion cells and still relayed action potentials along axons to the brainstem. Analyses of auditory responses of intact turtles would be useful to understand the central auditory effects of hypoxia (Christensen-Dalsgaard et al., 2012; Christensen-Dalsgaard et al., 2010; Patterson, 1966; Piniak et al., 2016; Willis, 2016)

### Visual responses measured ocularly and centrally are both sensitive to hypoxia

The amplitudes of both ERGs and OT field potentials were decreased when superfused with near-anoxic media, a predictable outcome given that phototransduction is mediated by an energetically expensive enzymatic cascade found in the retinal photoreceptors. The retina, in general, and the photoreceptors, in particular, use the highest amount of metabolic energy of all the vertebrate sensory systems (Country, 2017).

Experiments of retinal cellular physiology indicate that P_O2_ in the outer retina is most modulated by retinal illumination at the photoreceptor layer (Linsenmeier, 1986), more so in cones as compared to rods (Ingram et al., 2020). Presumably, the highest oxygen need would be in animals with cone-dominated retinae. The aquatic turtle should therefore be very sensitive to retinal anoxia, especially considering its cone-dominated retina and its lack of a blood supply to the outer retina (Damsgaard et al., 2019).

This study of anoxic turtles was consistent with an earlier ERG study of anaesthetized, nitrogen-ventilated turtles (Stensløkken et al., 2008). Although those *in vivo* ERGs were recorded using 10-fold longer light flashes during two-hours of anoxia, those ERGs were qualitatively similar to that of our preparation in that the b-waves in both studies were significantly diminished by the anoxia. In vivo a-waves were significantly decreased but their responses were smaller, like the a-waves in our study that tended to decrease, but also did not achieve statistical significance. Other reports found that a-wave responses in both hypoxia-intolerant and hypoxia-tolerant vertebrates are less sensitive to hypoxia than b-wave responses and central visual responses (Johansson et al., 1997a; Niemeyer, 1975; Tinjust et al., 2002a). From these comparisons, we conclude that the responses to anoxia of the b-wave observed *in vivo* is intrinsic to the eyecup, and likely reflect some photoreceptor component due to the phototransduction cascade that occurs within its outer segments.

### Differential Effects of Severe Hypoxia on the Components of the Visual System

When compared to light responses measured in the retina, the effects of severe hypoxia on responses recorded in the OT were quantitatively greater. This difference might be explained simply if hypoxia effects within the cranium occur independent from the response depression within the retina, and both ocular and midbrain effects are additive.

Another explanation for measuring strong hypoxic effect in the OT is that these two in vitro preparations are very different due to the experimental recording techniques, differences in the forms of response quantification and due to the biological tissues themselves. ERGs represent averaged, low frequency trans-retinal currents, as compared to the tectal field-potentials, derived from multi-unit spiking neurons. Those responses were also quantified using different physiological measures. Another difference is that ERGs sum the entire retinal surface, whereas the OT electrode receives just input from a localized region of the visual space that is mapped onto the tectal surface. Moreover, the neural architecture of these two visual structures differ substantially: retinal neurons relay graded potentials along the narrow radial path while tectal neurons gather synaptic inputs from have large complex dendritic trees that project laterally across the entire tectal surface.

A common feature of ERGs and OT field potentials is that their light responses were decreased during anoxic media. Light responses during hypoxia are also decreased in mammals including humans. Using isolated rabbit retina in vitro, total anoxia blocked all retinal light responses (A. Ames and Gurian, 1963). Using intact mammals, severe hypoxia or total anoxia was not feasible (but see (Eysel, 1978)). Hypoxic cats showed differential effects on the a-wave versus b-wave response, depending on the level of hypoxia (Derwent and Linsenmeier, 2000), see also (Noell and Chinn, 1950). Mild hypoxia (PO2 of 50–60 torr) reduced the a-wave amplitude by small amounts, while an 8.9% occurred due to severe hypoxia (20–30 torr). The same severe hypoxia had a strong (35%) effect on the b-wave amplitude. Humans breathing mild hypoxic air (12% O_2_ / 88% N_2_) exhibited a statistically significant decrease in their b-wave amplitude but no change in their a-wave amplitude (Tinjust et al., 2002b). So, like the turtle ERG, the mammalian a-waves due to photoreceptors are more resistant to hypoxia than b-wave activity of the inner retina.

The ability of the anoxia-tolerant crucian carp to respond to visual stimuli was also investigated in an anoxic environment. Unlike pond turtles, crucian carp are not quiescent when exposed to anoxic conditions (Nilsson et al., 1993), which suggest that their visual system may be better able to respond to light during anoxia, unlike pond turtles. Yet, ERGs and OT field potentials of the crucian carp had a strong decrease in visual responses (Johansson et al., 1997b). However, the rate of the response amplitude decrease in fish were slower than the decrease rates observed in the turtles, perhaps due to the unique fish eye (Country, 2017) or the ability of the *in vitro* turtle preparation to reach anoxic levels quickly by direct perfusion.

### Perspectives

These studies find that visual responses were significantly decreased in both retina and OT during hypoxic conditions while responses to auditory stimuli (direct vibration of the tympanic membrane) remained in severely hypoxic conditions (Po_2_ < 20 torr). The physiological mechanisms that underlie a turtle’s auditory capacity to tolerate hypoxia/anoxia may provide valuable insights into an organism subjected to anoxia: how animals manage with decreased ATP availability. Their survival usually requires organ-specific metabolic suppression (Hochachka, 1986) and physiological functions are likely prioritized, with less essential and more energetically expensive ones suppressed, and more essential and energetically inexpensive ones preserved (Jackson, 1987). Sensory functions pose a particularly compelling problem because their relatively high energetic cost will impact on the animal’s survival. Studying sensory function in anoxia-tolerant animals like turtles may enable us to understand where the different sensory functions sit on this spectrum, while also revealing fundamental mechanisms of sensory function in response to low oxygen.

The intact pond turtle as a model organism to observe recovery from strong hypoxia as it has been shown that the intact animal can survive extensive periods of time of anoxia during near-zero temperatures. Freshwater turtles whose winters are spent in sub-zero °C environments enter the nearby ponds to survive their winters. Surviving in anaerobic setting requires many biological adaptations that are only successful in near-freezing temperatures. These *in vitro* experiments were limited to controlling the Po_2_ directly in the physiological solution, yet not reducing body temperature. Obviously, the survival of an intact turtle in its natural environment may also be dependent upon other factors: temperature, blood chemistry, local pH in the brain and other systemic circulating modulators. Future *in vitro* studies should evaluate the effects of lower temperatures or other systemic parameters in normoxic media, in combination with hypoxic media.

## Acknowledgements

Supported by an NSF CAREER award (1253939) to DEW; ERG experiments were performed in partial fulfilment of Master’s Thesis (SA).

